# Acute and chronic physical activity improve spatial pattern separation in humans

**DOI:** 10.1101/2021.12.13.472386

**Authors:** Daniela Ramirez Butavand, María F. Rodríguez, María V. Cifuentes, Magdalena Miranda, Cristian García Bauza, Pedro Bekinschtein, Fabricio Ballarini

## Abstract

Physical activity benefits both fitness and cognition. However, its effect on long-term memory is unclear. Successful memory involves not only remembering information over time but also keeping memories distinct and less confusing. The ability to separate similar experiences into distinct memories is one of the main features of episodic memory. In this work, we evaluated the effect of acute and chronic physical activity on a new task to assess spatial pattern separation in a 3D virtual reality environment. We manipulated the load of memory similarity and found that 25 minutes of cycling after encoding - but not before retrieval - was sufficient to improve similar, but not dissimilar memories, 24 hours after encoding. Furthermore, we found that participants who engaged in regular physical activity, but not sedentary subjects, showed memory for the similar condition the next day. Thus, physical activity could be a simple way to improve discrimination of spatial memories in humans.

---

Physical activity (PA) has been shown to benefit both fitness and cognition. In particular, Blondell and colleagues (1) have shown that regular aerobic exercise protects against cognitive decline. Also, regular aerobic exercise considerably improves the performance of tasks that involve attention, processing speed, and executive function (2). However, the relationship between aerobic exercise and memory is not well understood. Voss and colleagues (3) have summarized the findings linking PA and memory and concluded that there is a positive effect of aerobic exercise on different memory tasks such as visuospatial, story recall, and relational either for cognitively normal middle-aged or for older adults. On the other hand, Roig and colleagues (4) concluded in their review that long-term cardiovascular exercise has no significant effect on long-term memory and produces only small improvements in short-term memory. Interestingly, no negative effects of PA were found on cognitive functions. Even though most of the above-mentioned studies have shown numerous benefits of aerobic exercise, most of them have evaluated these benefits on a chronic basis. Since most people face complex and highly demanding routines that usually jeopardize their compromise in maintaining long-term exercise routines, these benefits do not apply to most cases. Several studies have shown that the rate of abandonment of these practices is very high and is due to various factors mainly related to lack of time (5). Therefore, differentiating the effects of chronic from acute physical activity is highly relevant to the clinical application.

Other studies have focused on the effect of short bouts of exercise on various cognitive functions. A single session - of variable duration - of moderate aerobic exercise has been shown to improve numerous executive functions such as attention (6), inhibitory control (7), and cognitive flexibility (8), among others (for a review, see (9)). In addition, several investigations studied the effect on short-term memory, showing a positive effect of a 10-minute walk followed by a 15 to 30-minute rest on the recall of words learned immediately after exercise (10). Also, they showed that 30 minutes of cycling before learning a story improved the short-term retention assessed at 30 minutes (11) compared to the control group. Although these studies improved our understanding of the acute effects of exercise on memory formation, few studies evaluated the impact of exercise on long-term memory. The main studies in this field found associated improvements in the consolidation of motor, emotional, and picture-location memory (12–14). However, it is still necessary to keep studying the effect of these ecological interventions on different phases and aspects of memory.

In particular, in this research, we focus on studying the effect of exercise on the pattern separation process, which is a critical feature of episodic memory (15). Computational models have defined ‘pattern separation’ as the process of orthogonalizing two similar inputs into distinct outputs. This ability deteriorates with aging and Alzheimer’s disease (16– 18), so it is essential to study simple ways to improve it. Behaviorally, pattern separation has been studied using different tasks in both rodents and humans (19–21). Rodent studies have established an important role for the dentate gyrus (DG) of the hippocampus in spatial pattern separation (22). However, the tasks used in humans mainly use objects as to-be-remembered stimuli. In addition, regular and acute PA seems to improve this ability to differentiate memory in humans (23, 24). However, no studies have evaluated the effect of exercise on the disambiguation of long-term memories. Furthermore, no study in humans has addressed the effect of PA directly on spatial ‘pattern-separated’ memories.

In this study, we evaluated the effect of acute and chronic physical activity in a new task to assess spatial pattern separation that is similar to that used in rodents (20), which allows parametric manipulation of memory similarity during encoding. The activity was carried out with the support of a software solution, including a post hoc virtual reality environment (Computer Assisted Virtual Environment, CAVE), which provides a greater sense of immersion than tasks performed with computer screens (25, 26), and, at the same time, allows the participant to navigate the spatial context of the study in the same way that rodents do. Based on the knowledge of the molecular mechanisms that subserve both these types of memories and PA in rodents (22, 27–29), we hypothesize that an acute bout of exercise during the consolidation of similar spatial memories may improve long-term retention. Additionally, people who engage in frequent physical activity may benefit from an increase in memory retention of similar positions.

## Results

To investigate the effect of PA on the discrimination of similar spatial memories, we recruited sedentary subjects for the control and acute groups in both similar (20°) and dissimilar (40°) conditions and athletes for the chronic group. This distinction was based on the participants’ IPAQ (International Physical Activity Questionnaire) reports, using only the questions related to leisure activities. The chronic group had a significantly higher score compared to the control and acute groups, and no differences were observed between the latter two groups (Kruskal-Wallis analysis, 20°: H = 19.22, *p* < 0.0001 and 40°: H = 12.59, *p* = 0.0018).

In the first part of the protocol, the participants performed the pre-TR phase. After the second trial, they were instructed to use spatial strategies to orient themselves in the arena. Accuracy was measured as the difference between the target and the selected flag positions (distance to target). A repeated-measures ANOVA analysis revealed a significant effect of trials (20°: *F*_(4,32)_ = 14.64, *p* < 0.001, *η*^2^ = 0.28; 40°: *F*_(4,32)_ = 20.63, *p* < 0.001, *η*^2^ = 0.22) and there were neither differences between groups (20°: *F*_(2,16)_ = 0.03, *p* = 0.967; 40°: *F*_(2,16)_ = 0.06, *p* = 0.939) nor interactions (20°: *F*_(8,64)_ = 1.98, *p* = 0.064; 40°: *F*_(8,64)_ = 0.82, *p* = 0.591). This indicates that participants improved their performance throughout trials.

The accuracy in the control test (using a separation between flags of 60°) was similar for all groups (ANOVA test: *F*_(5,50)_ = 1.61, *p* = 0.18), indicating that all groups began with a uniform ability to use the joystick and a similar spatial orientation in the CAVE.

Each participant was assigned to one of the two conditions (*a* = 20° or *a* = 40°) and performed the TR1 that was evaluated immediately afterward (delay 0h). We calculated the absolute value of the difference between the flag positioned in the middle of the two flags that appeared in the TR (target) and the selected flag. No differences were observed between groups in the two conditions (20°: *F*_(2,26)_ = 0.42, *p* = 0.66; 40°: *F*_(2,24)_ = 0.94, *p* = 0.41).

To find out whether exercise influenced the consolidation of similar memories, participants performed a second training session (TR2) followed by an intervention (cycling in the acute group or watching a video in the control and chronic groups) that was tested 24 hours later (TS2, delay 24h, Figure 1**a**). The mean distance covered by the participants on the stationary bike was 7.29 ±0.86 *km* in the 20° group and 6.88 ±0.70 *km* in the 40° group (no differences were found between groups, *t* = 0.90, *p* = 0.39). The distance to the target measured 24 hours after learning was significantly different between the groups in the similar condition (Figure 1**b**, 20°: *F*_(2,26)_ = 4.95, *p* = 0.015, *η*^2^ = 0.28). Post hoc pairwise comparisons showed that the acute group could solve the task better, obtaining smaller distances to the target than the control group (*p* = 0.012, d = 1.43). This outcome was not observed in the dissimilar condition. The physical intervention had no impact on the group that solved the task with a greater separation between flags (Figure 1**c**, 40°: *F*_(2,24)_ = 0.06, *p* = 0.94). We analyzed performances separately in male/female and the 2-way ANOVA did not reveal significant differences between sexes (20°: *F*_(1,23)_ = 0.48, *p* = 0.50; 40°: *F*_(1,21)_ = 0.37, *p* = 0.55).

**Fig. 1.**
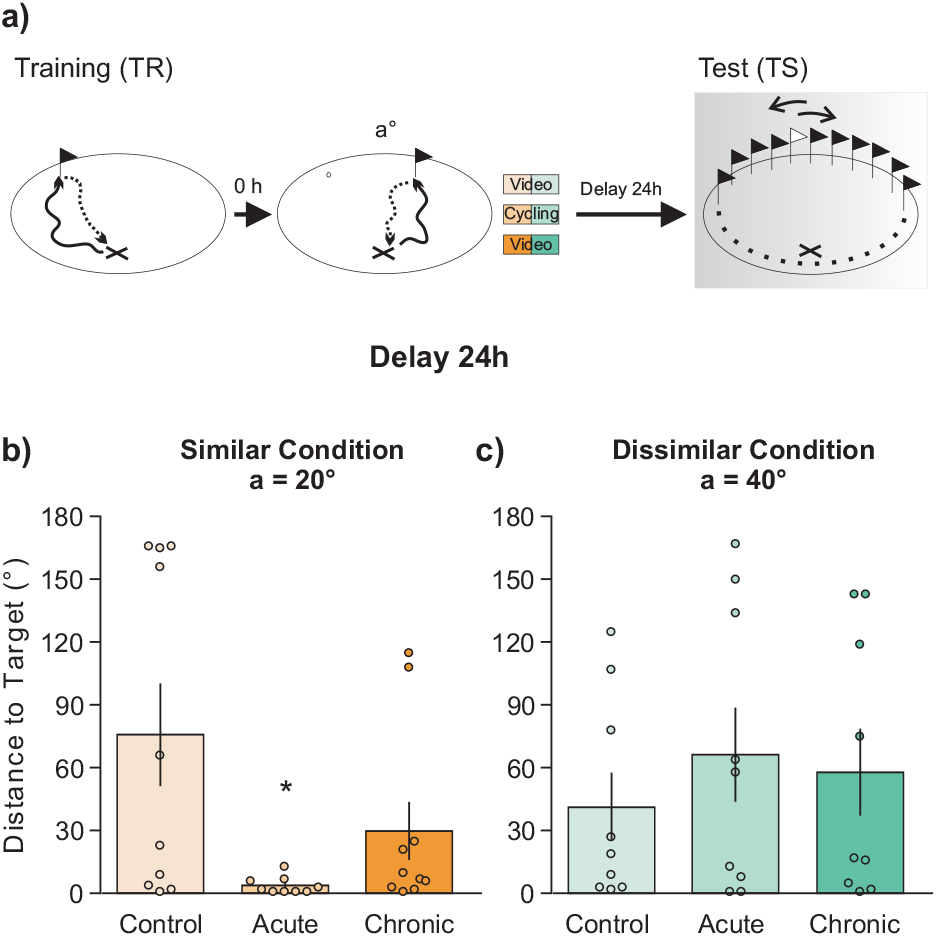
Effect of exercise on the consolidation of similar and dissimilar memories. **(a)** Schematic representation of the training and testing phases for the delay 24h. The performances were calculated as the distances to the target (the position of the flag in the middle of the two flags learned in the TR minus the position of the flag selected by each participant in the TS) and are shown as the mean ±SEM for the different groups that performed the task using **(b)** an angle of 20° or **(c)** an angle of 40°. One-way ANOVA revealed significant differences only in the 20° condition with a 24h-delay *p* < 0.05. The number of participants in each group, from left to right, was 10, 9, 10, 9, 9, and 9.

Afterward, participants carried out the last training (TR3) which was tested immediately (the responses of two participants in the 20°-chronic group were lost). Similar performances were observed for all groups under both conditions (20°: *F*_(2,24)_ = 1.03, *p* = 0.37; 40°: *F*_(2,24)_ = 2.58, *p* = 0.097).

We reasoned that if PA allowed for long-term memory, both the acute and chronic groups would not select the flag randomly during the long-term memory test (TS2). Thus, we analyzed the distributions of the data obtained to determine if participants were choosing a flag at random leading to a uniform distribution among all possible positions (i.e. ‘no memory’) or if they remembered the positions of the flag where the data distribution should be around zero. We did this by comparing the performance distributions with 1000 uniform distributions randomly generated using a Kolmogorov-Smirnov test and then we averaged the *p*-value obtained in each iteration. Results indicated that all distributions of delay 0h were significantly different from the random uniform distribution (*p* < 0.05). In other words, all groups were able to remember the locations of the flags immediately after TR. In contrast, for the 24h delay, we observed that all distributions were similar to the uniform distribution, except for that of the acute and chronic groups in the 20° condition (Figure 2**a**, *p* < 0.01 and *p* < 0.05, respectively). This may reflect that these groups did remember the position of the flags learned the previous day. Conversely, the control group in the similar condition and all groups in the dissimilar one (40°, Figure 2**b**) showed no memory about the position of the flags (*p* > 0.05 in all cases).

**Fig. 2.**
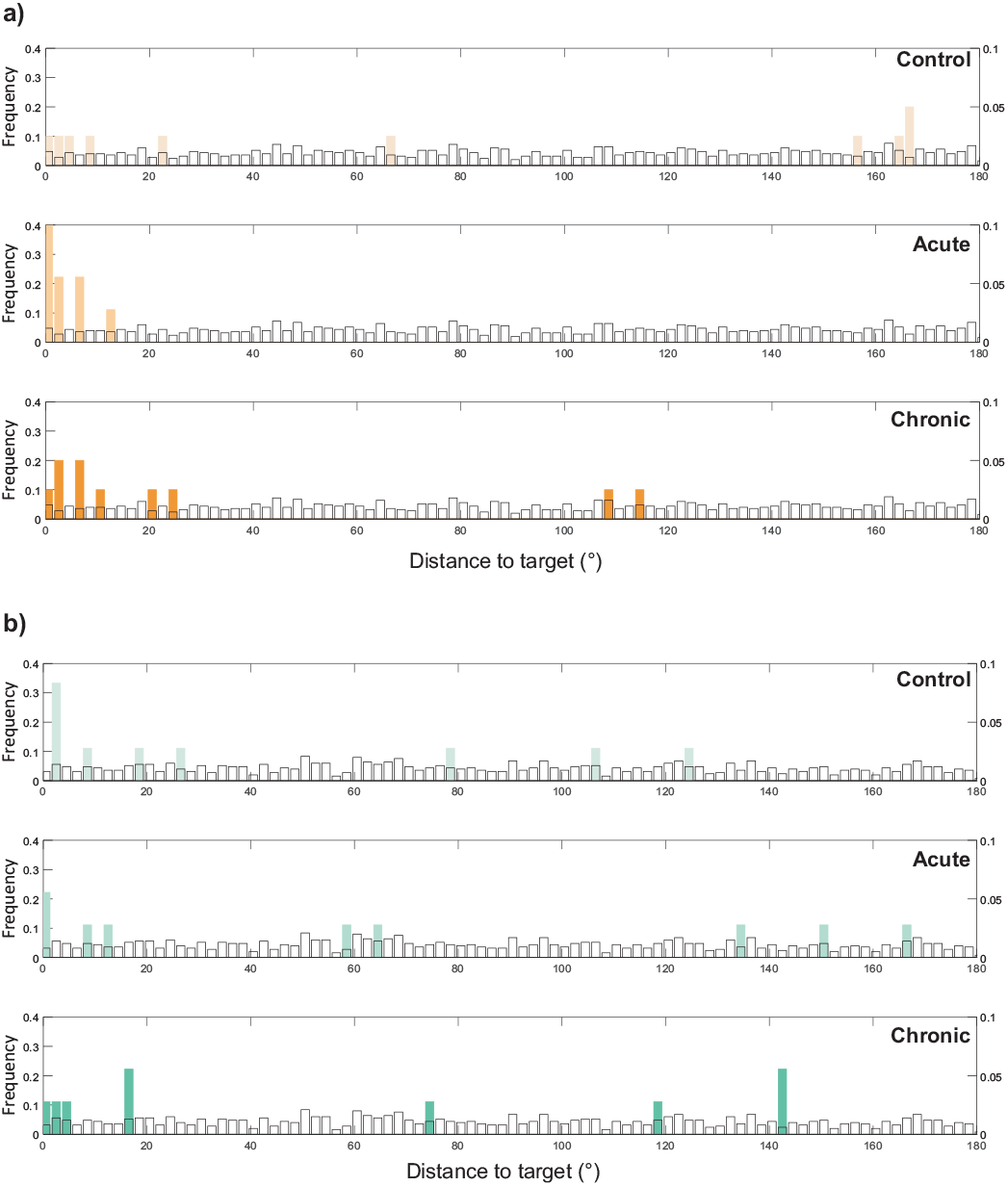
Comparison between performance distribution and uniform distribution. We plotted the frequency obtained for each possible angle (i.e., how many participants obtained a particular distance to the target). The orange bars represent the participants who were in the similar condition (20°) **(a)** and the green bars in the dissimilar condition (40°) **(b)**. Also, we plotted a uniform distribution generated randomly (white bars). The uniform distribution plotted is just representative because the distributions used in the comparison were of the same length as the data. We compared the performance distribution with the uniform distributions in each case using a Kolmogorov-Smirnov test. We obtained significant differences only in the Acute and Chronic groups in the 20° condition: *p* < 0.01 and *p* < 0.05, respectively.

Finally, to investigate whether exercise additionally modulated retrieval of similar spatial memories, we recruited a different group of participants to perform a similar protocol but, instead of cycling or watching the video immediately after TR2, they spent 25 minutes cycling on a stationary bike or watching a video just before the TS2, when they were asked to remember the position of the flags learned the day before (Figure 3**a**). No differences were observed (*p* > 0.05) between the control and the acute groups on either of the two delays: 0 and 24 hours (Figure 3**b** and 3**c**, respectively). Together, these results suggest that performing physical exercise immediately after encoding - but not before retrieval - promoted the long-term retention of similar spatial memories, with no effect on memories that do not require pattern separation.

**Fig. 3.**
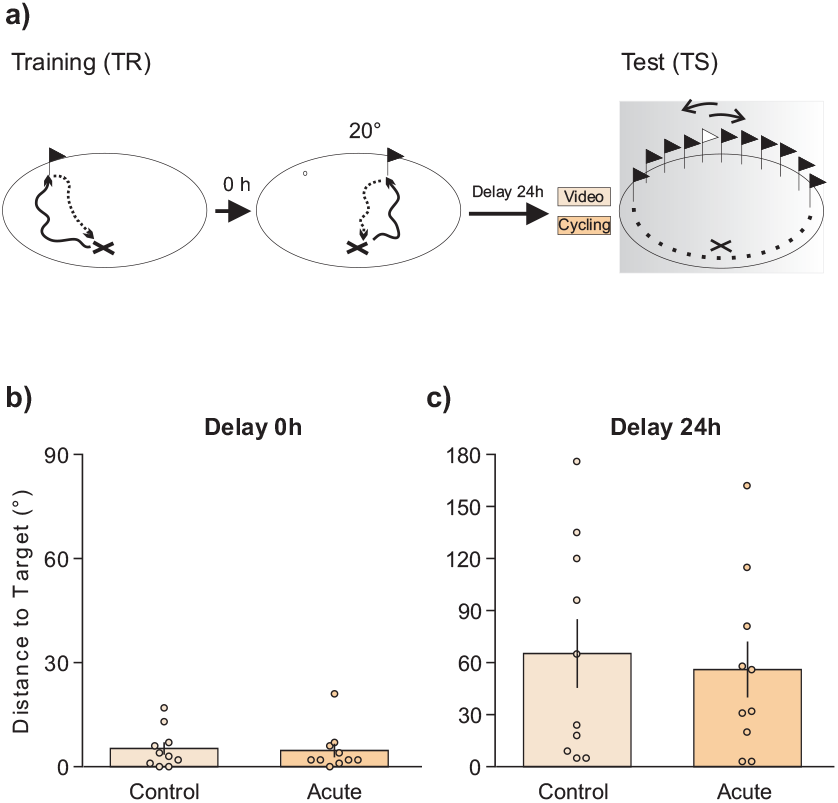
No effect of exercise on the retrieval of similar memories. **(a)** Schematic representation of the training and testing phases. Performance was calculated as the distances to the target (the position of the flag in the middle of the two flags learned in the TR minus the position of the flag selected by each participant in the TS) and is shown as the mean ±SEM for the control and acute groups. Both groups performed the task using a 20° angle for both: the delay of 0h **(b)** and the delay of 24h **(c)**. The acute group exercised before retrieval. *t* -test did not reveal significant differences in either of the two conditions *p* > 0.05. The number of participants for the control and acute groups was 10 and 10, respectively.

## Discusion

In this work, we developed a task to evaluate spatial pattern separation in humans and studied the effect of acute and chronic physical activity on both the consolidation and the retrieval of overlapping spatial memories. The results showed that a single bout of acute physical activity after encoding, but not before retrieval, was sufficient to improve the long-term retention of similar, but not dissimilar, spatial memories. Additionally, both the participants in the acute group and those who engaged in regular physical activity selected the position of the flag in a non-random way.

The task developed here allowed the participant to interact with and navigate through a virtual reality environment. The participants could visualize the target in a spatial context and then be able to recall its position relative to the background closer to a naturalistic manner. In addition, the virtual reality device allowed the exact reproduction of the conditions between each participant, without depending on the exploration of environmental or climatological events. Previous studies have shown that, on the one hand, spatial information can be reliably transferred between real and virtual reality environments when the virtual reality environment was modeled from the real (30), and, on the other hand, that the brain areas involved in both scans are analogous (31–33). Therefore, developing tasks that allow evaluating different cognitive abilities in virtual environments provide valuable tools that allow greater precision when navigating spatial contexts. Another advantage of this task is its analogy to the one previously used in rodents (20), something that stands out for the unprecedented benefit of its ability to nurture from the growing molecular understanding that this field has achieved in animal models. The most frequently used task to evaluate pattern separation in humans was developed by Stark’s laboratory (21). In that task, the participant must distinguish similar objects of daily life immediately after encoding. Given the role ascribed to the hippocampus in spatial memories (34, 35) and, specifically in pattern separation (22), it is curious that few investigations have studied pattern separation in spatial types of memories. We believe this is relevant given the present debate on whether the DG of the hippocampus is a general pattern separator or it is only important for spatial aspects of memories (36). The only spatial task that has been used so far consists of the appearance of a point on the computer screen during the training. During the testing phase, immediately after training, the participant must select the position of the learned point between 2 options (37). However, the lack of context makes this type of spatial task less ecological and distant from real everyday experiences. Conversely, we have recently demonstrated that to correctly solve our pre-training task within the CAVE, it is necessary to use spatial strategies (38). In addition, those tasks do not modify the similarity of the memories during encoding, when pattern separation is proposed to occur (39).

Although there is consensus in the scientific community that physical exercise benefits memory, it is not clear the type of exercise, its duration, and/or timing. Moreover, most of the effects related to memory enhancement have been found in typical laboratory memory tasks, making it difficult to predict whether everyday memories would be improved in the same way. Our results demonstrated that 25 minutes of exercise improved the consolidation of learned information within the virtual reality immersive environment only when the position of the learned flags were close to one another, but not when they were more distant. This means that this improvement is specific for spatial memories that require disambiguation due to their similarity and not for memories that are already easily distinguishable. Moreover, the exercise did not have a significant impact on retrieval. These results suggest the existence of specific time windows of exercise effectiveness (after learning and not before retrieval). No study in humans, to our knowledge, has evaluated the effect of physical activity on the different phases of long-term memory of similar and dissimilar experiences. Therefore, it is relevant to continue the in-depth study of different interventions that may result in direct applications to the clinic. In addition, we observed that the distribution of performances of the chronic participants differed significantly from a uniform distribution, indicating an improvement in memory differentiation also for participants who perform physical activity regularly. Therefore, we can conclude that both acute and chronic exercise showed benefits in this spatial differentiation task. However, it is worth pointing out that the benefits of physical activity in this task appeared to be more effective for the acute than for the chronic group. We acknowledge that the studies did not involve large sample sizes, but given the consistent pattern and large size effects, we think our hypothesis has been effectively supported. There are numerous studies where differences were found between men and women in terms of spatial strategies or cognition, both in real (40) and virtual reality environments (41). However, in this work, we did not find differences due to sex. This could be due to the small sample size, although recent work supports our results (30).

Previous studies have demonstrated that a single bout of PA immediately after learning was also able to improve the long-term retention in motor and emotional memories (12, 13). Furthermore, van Dogen and colleagues (14) have shown that a single bout of PA immediately after learning had no effect on a picture-location memory but did have an effect when it was given four hours after learning. Therefore, the time window of exercise effectiveness could be different for the different types of memories studied and the similarity of the to-be-remembered items.

In addition, Borota and colleagues (42) have shown that caffeine administration after learning - and not before retrieval - improved the long-term memory of the task evaluating the recognition of similar and dissimilar daily life objects during retrieval. So, they showed a time window similar to the one we found in which similar memories could be enhanced. In fact, the authors suggest that caffeine administration could be increasing Brain-Derived Neurotrophic Factor (BDNF) levels (43) and thus improving memory.

Our results are consistent with previous studies in rodents addressing the role of BDNF in the analogous task (22). In these experiments, blockade of BDNF impaired the consolidation of similar but not dissimilar memories and had no effect on retrieval. Although the mechanisms that are mediating exercise-induced similar memory enhancement were not studied in the present study, one possibility is that exercise increases BDNF expression, leading to an enhanced memory differentiation. In this sense, it has been shown that both acute and chronic exercise induce BDNF expression (28, 44) in rodents in the DG, a region implicated in the discrimination of spatial memories (22, 45). Furthermore, in humans, serum BDNF levels increase with exercise, mainly after an acute bout of exercise ((6); for review see: (46, 47)). On the other hand, given that BDNF activity in DG is necessary for the consolidation of similar spatial memories (22), the 25-minutes cycling could provide the BDNF needed to consolidate similar flag positions. Moreover, in rodents, exogenous BDNF injected in the DG immediately after training increased discrimination of spatial memories, similar to what we found after a short period of PA.

Another possibility is that exercise is acting through an increase in the levels of neurogenesis: several studies have demonstrated that the DG - and in particular its immature neurons - plays a fundamental role in pattern separation (19, 48, 49). Furthermore, it has been shown that aerobic exercise increases neurogenesis in the DG (50, 51). However, this mechanism could not explain the results obtained for the acute group since the time windows of cell proliferation and maturation (weeks) and the BDNF requirement for consolidation of similar memories (minutes to hours) do not coincide.

In a recently published opinion (52), Quian Quiroga argued that there is no pattern separation in humans, however, our results provide evidence to this debate in the opposite direction, since an improvement was seen specifically when the positions of the flags were similar, which supports the idea of different processing of similar and dissimilar experiences.

Altogether, our findings provide evidence for the efficacy of exercise interventions as a valuable tool for increasing declarative memory consolidation in humans, specifically when the encoding phase presents similar spatial locations. The low cost, healthy and practical nature of exercise makes it ideal for interventions in educational and clinical settings. Our experiment thus serves as a proof of principle study that could inspire future applications of exercise to boost long-term episodic memory in various populations.

## Materials and Methods

### Participants

Seventy-six healthy young adults (25 women) participated in this study. Of the total number of participants, 19 were athletes (5 women), recruited from the athletic group of the Universidad del Centro de la Provincia de Buenos Aires. The remaining 57 participants belonged to the sedentary group, who declared not to have done physical exercise over the last year. The mean measured body mass index (BMI) of the sample was 24.26 ± 4.58 *kg/m*^2^ (no differences were found among groups). All participants had normal or corrected-to-normal vision and normal color vision. The inclusion criteria required that all participants were between 18 and 35 years old (24.11 ± 4.68), did not consume psychotropic drugs, were native Spanish speakers, did not report neurological or neuropsychiatric diseases. All subjects were required to refrain from consumption of caffeine during the training and testing day.

This study was reviewed and approved by the Ethical Committee Instituto de Tisioneumonología “Prof. Dr. Raúl Vaccarezza”. Written informed consent to participate in this study was provided by the participants.

### Software Solution

We have developed our spatial task using a software toolbox that we created based on similar works (53, 54). We developed a Virtual Environment Editor to design the virtual arena, a Protocol Editor to create the task, and a Data Collector Module to save information relevant to the study. These three components work together to build the task. They are connected to a virtual reality device to visualize and perform the task (see: (38)).

With the Virtual Environment Editor, we created and configured the virtual scene where the experiments were carried out. In a virtual reality environment, we can simulate from an open place such as a valley, a forest, a navigation in the ocean to a closed environment such as the interior of a house, a room, the facilities of a building, the rooms of a factory, among others. In this work, we used the image of flat sand with some geographical features with which the subject could orient him/herself.

Protocol Editor is a simple graphical user interface to build tasks. It is composed of an editor of diagrams of transition states to design different tasks. With Protocol Editor we modeled the states of the participant during the task. The participant is linked to a particular task instruction at each time. Protocol Editor processes the commands executed by the user, communicates the virtual reality environment and keeps tasks synchronized.

Data Collector Module saves information recorded by virtual reality devices.

As a virtual reality device, we used a CAVE. CAVE is a room-sized cube in which the walls and floor are projection screens generating the feeling of immersion of the user (55). Four to six projectors are used to display virtual reality environments on the walls and floor. Each of the projectors is connected to a dedicated PC responsible for generating the image. In the CAVE the subject can move, giving the feeling of total immersion (26) and creating the illusion of having completely unlimited space.

We used the Rubika CAVE, which was developed by members of the PLADEMA Institute (Argentina) in 2014. It consists of a three-meter cube-shaped steel structure. Its inner walls are covered with special fabrics to allow the rear-projection. The projectors used to display the images onto the walls have a 1280×800 resolution, while the one projecting the floor image has a 1980×1080 resolution. Furthermore, four PCs hosted the applications in charge of managing each projection including the effects of the user’s actions.

A joystick allows users to move inside the CAVE and interact in different ways within the application. The participant can move forward, backward, or sideways and could also rotate in the spot using the joystick.

With our software solution, a wide range of spatial tasks can be created and performed in several virtual reality devices. The software has been made available in a permanent third-party archive (https://osf.io/jbfpn/). In this OSF project, we include a wiki and videos with information and instructions for using our software. We give a desktop version of a CAVE, but we encourage interested parties to use it on other virtual devices. In the OSF project, we explain how to configure our Behavioral Spatial Pattern Separation Task.

### Behavioral Spatial Pattern Separation Task

We designed a task, similar to that used in rodents (20). The protocol consisted of three stages: pre-training, training, and test.

During the **pre-training** (pre-TR) phase (38), the participant was located in the center of the arena with a background consisting of a fairly uniform landscape with some valleys and mountains. At first, a flag appeared in one of the 360 possible random positions within an imaginary circumference of 20 *m* radius. The subject walked through the virtual arena with the joystick, picked up the flag, and then returned to the mark in the center. Once he/she arrived, a total of 360 flags appeared and the subject was asked to select the exact flag position that was previously collected, using a cursor. Immediately after, the placemark changed its position and the participant walked to the new starting point. At that moment, the second trial began with the appearance of a new flag, and the procedure was repeated. The number of trials per participant was variable. Trials were finalized after i) the participant selected the correct distal flag three consecutive times or ii) the procedure was repeated ten times, whichever came first. For each trial, we calculated the distance between the selected and target flags with our software solution.

During the **training** (TR) phase, the participant was located once again in the center of the arena and a new flag of a different color appeared. The subject picked it up and returned to the starting point. Immediately after, a second - identical - flag appeared in a new position separated from the previous flag position by an angle of *a* = 20° or *a* = 40° within the same circumference (Figure 1**a**, left). This phase was finalized once the participant collected it and returned to the initial mark.

During the **testing** (TS) phase, 360 flags appeared, on the same circumference, and the participant had to select the flag that was positioned exactly in the middle of the two flags collected in the training phase (Figure 1**a**, right).

We created and configured the three phases of the task using the Protocol Editor of our software solution. In the third-party link, we describe how we did it: https://osf.io/fn9r8/wiki. The rationale for the task was that if the participants were able to form separate representations of the two nearby positions seen during training, the representations of the two locations should be distinct and resistant to confusion; therefore, they should be successful at distinguishing where the center was. However, if the representations of the two locations were not sufficiently separated, the position of the center would be chosen randomly.

The protocol performed by the participants started with the pre-TR phase, during which the participant got used to the virtual reality environment and generated a spatial map of the arena. Following this, all participants performed a TR with a spacing angle between flags of 60° and were asked to select the right-handed flag (control test). The goal of this test was to assess the accuracy in solving the task before starting the relevant tests. Afterward, participants were randomly assigned to one of the following conditions: 20° (Similar Condition) and 40° (Dissimilar Condition). All subsequent training was done at the same angle. Following the control test, participants carried out the first training (TR1) and immediately after, the first test TS1 (delay 0h). After completing TS1, they watched a video (to avoid any interference between trials) for half an hour and began the second training (TR2). Once finished TR2, they were randomly assigned to the control, acute, or chronic group. The control group was made up of sedentary people who watched a video about bicycle races for 25 minutes (similar to previous studies: (13, 56)). The acute group was also made up of sedentary people but who, instead of watching a video, cycled for 25 minutes. Finally, the chronic group was made up of athletes who watched the same video as the control group. After this, the first day ended and the participants returned 24 hours later to perform the second test (TS2) corresponding to the learning of the previous day (delay 24h). Lastly, they carried out the last training and test (TR3 and TS3, respectively) to assess the specificity of the physical intervention. Before leaving the laboratory, participants filled in a short version of the International Physical Activity Questionnaire (IPAQ), which was used as a measure of self-reported physical activity. Participants were asked to report the amount of walking, and the number of times they had performed moderate and vigorous activities over the previous seven days.

With an independent group of sedentary subjects, we evaluated the effect of the intervention of physical activity on the retrieval of this memory. To do this, the subjects performed the same protocol but instead of separating the experimental groups after TR2, they watched the video on bicycle races (control) or cycled (acute) just before TS2.

### Physical Activity

A 25 min cycling task served as the acute exercise intervention carried out on a stationary bike. It started with a 5 min warm-up at the first resistance, after which the resistance augmented at the second level. They were asked to maintain the speed constant between 15 *™* 17 *km/h*, whichever was most comfortable for them.

### Statistical Analysis

Data are presented as the distance to the target (°) calculated with our software solution as the absolute value of the position of the central flag (in the middle of the two learned flags) minus the position of the flag selected by the participant. The results were presented as the mean ± SEM. When the data did not follow a normal distribution, it was transformed using the natural logarithmic function and then analyzed using the one-way analysis of variance (ANOVA). In cases of significant differences, post hoc analyses were made with the Tukey test when in one or both factors, the *p*-value was significant. A result was considered significant when *p* < 0.05. The data of the performance distributions were compared with 1000 uniform distributions randomly generated using a Kolmogorov-Smirnov test and then we averaged the p-value obtained in each iteration. All data were analyzed using GraphPad, Matlab, and Jasp software.

## ACKNOWLEDGMENTS

We thank Jorge Medina for his useful comments on the manuscript. This work was supported by research grants from the National Agency of Scientific and Technological Promotion of Argentina (ANPCyT) to P.B. (PICT 2012-1119, PICT 2015-0110 and PIP 0564).

